## Supplementary material for "Knockout of cyclin dependent kinases 8 and 19 leads to depletion of cyclin C and suppresses spermatogenesis and male fertility in mice": Suplementary Figures

### Slide 1
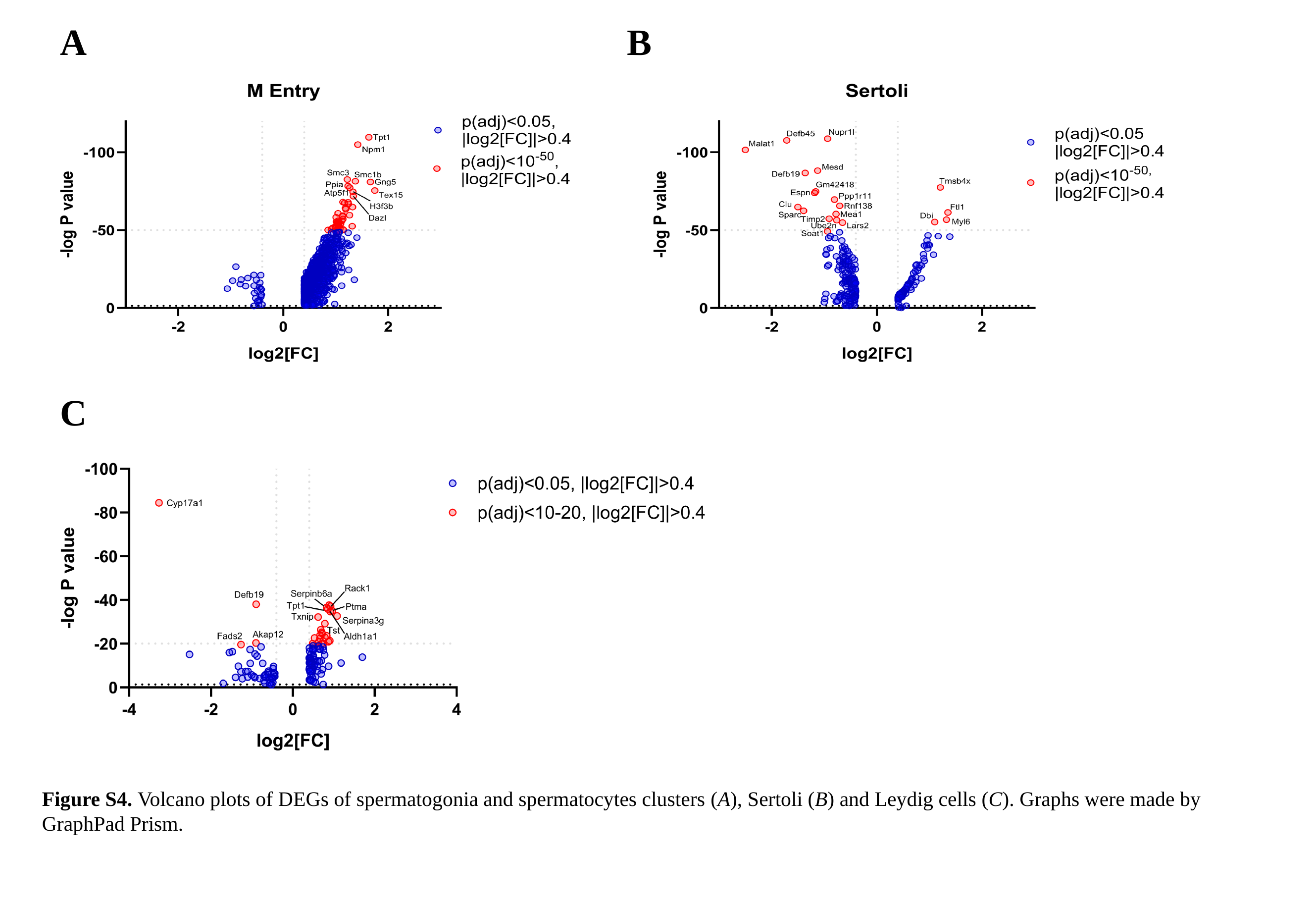

A
B
C
Figure S4. Volcano plots of DEGs of spermatogonia and spermatocytes clusters (А), Sertoli (B) and Leydig cells (C). Graphs were made by GraphPad Prism.
